# Synonymous mutations in AAV Rep enhance genome packaging in a library selection

**DOI:** 10.1101/2024.10.08.617207

**Authors:** Tasfia Azim, Dru Myerscough, Jonathan J. Silberg

**Author notes:** Jonathan J. Silberg, Department of Biosciences, Rice University, 6100 Main Street, MS-140, Houston, TX, 77005.

## Abstract

When producing Adeno-Associated Virus (AAV) gene therapies, a significant fraction of capsids can lack the desired DNA cargo. In AAV, Rep proteins mediate DNA packaging and virus assembly, suggesting that changes in Rep activity, expression, or DNA binding might affect genome packaging. To understand how mutations in the Rep gene affect activity, we selected a library of Rep mutants for their ability to produce active virions. By sequencing the Rep gene following the purification of viruses that package AAV genomes, we identified Rep mutants having non-synonymous mutations with a range of cellular activities. Surprisingly, synonymous mutations within the p19 promoter were enriched to the greatest extent, increasing in abundance by 10^2^ to 10^4^-fold. When the most highly enriched mutant was used to package a synthetic DNA cargo into the AAV capsid, the packaging efficiency could not be differentiated from native Rep. These findings suggest that these synonymous mutations enhance AAV genome packaging into capsids by affecting Rep-DNA interactions. They also suggest that silent sequence changes in the DNA cargo packaged by Rep can be used to tune packaging DNA packaging efficiency.

**Graphical Abstract:** 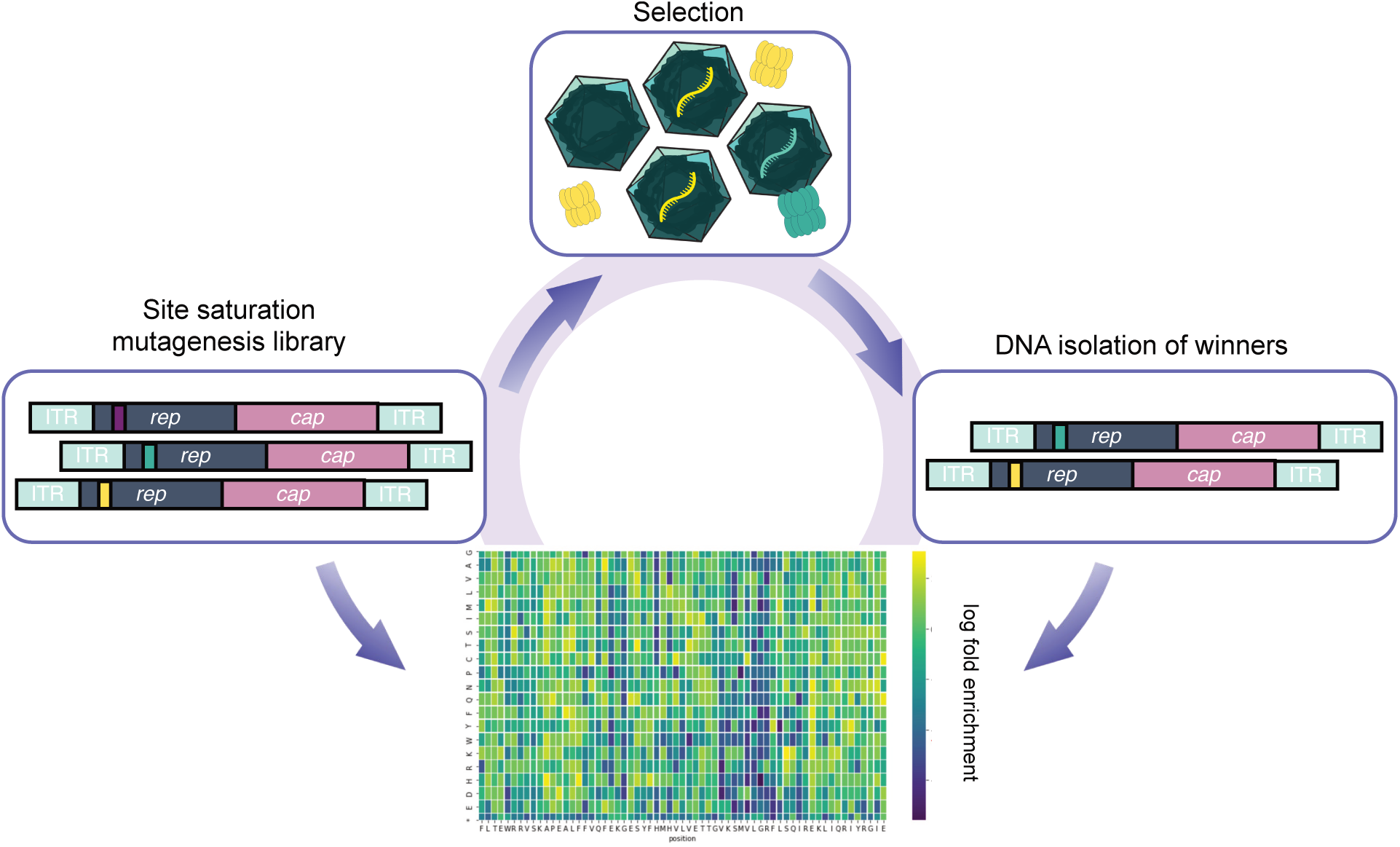

## Introduction

Adeno-associated virus (AAV) has become one of the most promising gene therapy vectors, with three AAV therapies federally approved for clinical use and numerous AAV therapies in clinical trials (1–3). However, manufacturing AAVs cheaply and efficiently remains a limitation. Raw materials for transient transfections to make AAVs, including plasmids and tissue culture reagents, are costly. Additionally, empty and partially filled AAV capsids are produced following transfections, with a spectrum of packaging levels reported across AAV preparations, ranging from 50 to 98% empty capsids (4–6). This heterogeneity in capsid packaging leads to multiple challenges. The presence of empty viral particles within preparations increases the gene therapy dosages required in clinical applications, which can cause undesired immune responses.

AAV is a small (25 nm), nonenveloped virus in the *parvoviridae* family (1). This virus is part of the *dependoparvovirus* genus, as it is incapable of replicating in the absence of a helper virus (1). The native capsid packages a 4.7-kb genome, which encodes nine proteins via alternative splicing and alternative start codon usage in two genes: *cap* and *rep* (7). These genes are flanked by hairpin DNA structures, which serve as packaging signals and start sites for genome replication. In the genome, the *rep* gene encodes four proteins (Rep78, Rep68, Rep52, and Rep40), while the *cap* gene encodes the three capsid proteins (VP1, VP2, and VP3), assembly-activating protein, and membrane-associated accessory protein (8, 9). To date, the capsid proteins have had their sequence-structure-function relationships studied to the greatest extent (10–14).

The Rep proteins form dynamic oligomeric structures that play several roles in the AAV lifecycle (15). Each Rep is made up of different combinations of the origin-binding domain (OBD), oligomerization domain (OD), and helicase domain (HD), and zinc-finger domain (ZFD) (15). All four Rep isoforms possess the HD and helicase activity but only Rep68 and Rep78 have an OBD. The OBD exhibits site-specific DNA-binding and endonuclease activities that play critical roles in DNA replication, transcriptional regulation, and site-specific integration into human chromosome 19 during latency (15, 16). Composed primarily of the HD, Rep52 and Rep40 facilitate DNA packaging into the capsid (17, 18). While structures of oligomeric Rep have been resolved in complex with ssDNA using cryo-EM (15), our understanding of this enzyme is not yet sufficient to rationally engineer it to maximize DNA packaging.

The dynamic and oligomeric nature of Rep makes it challenging to anticipate the effects of mutations on catalytic activity and interaction with DNA substrates. Recent work revealed that Rep proteins created by shuffling OBD from different serotypes can present higher packaging efficiencies than the parental Rep proteins recombined (6). In addition, a recent study analyzing the effect of amino acid substitutions on Rep activity identified mutations in multiple domains that increase DNA packaging (19). However, this study did not directly investigate the effects of synonymous mutations on Rep-mediated DNA packaging. Herein, we describe a similar combinatorial experiment wherein we evaluated the effects of mutations on Rep’s ability to package DNA. In this study, we sequenced the combinatorial diversity within the Rep gene directly, rather than using barcodes that are coupled to each mutation (19). This approach identified several synonymous mutations within the p19 promoter region increase Rep DNA packaging activity.

## Materials and Methods

### Plasmid constructs

All plasmids are listed in Table S1. The reporter plasmid (pAAV-CMV-GFP) was from Addgene, #67634 (20), while the helper plasmid (pXX6) was previously described (21). The parental pITR-p5-cap9-rep2 construct was built using Gibson Assembly (New England Biolabs) of gene block fragments of *rep* and *cap* purchased from Twist Biosciences (22). These genes were designed using AAV2 and AAV9 sequences, NCBI accession number J01901.1 and AX753250.1, respectively. This p5-rep2-cap9 fusion was cloned into a backbone with ITR sequences and then the fully assembled genome was cloned into the pAAV-CMV-GFP construct to create pITR-p5-rep2-cap9. To create pp5-rep2-cap9_K340H, a K340H Rep mutation was generated in pITR-p5-rep2-cap9 using PCR-based site-directed mutagenesis and primers (Integrated DNA Technologies). To create a positive control vector for selections (pITR-p5-rep2-cap9_bc), a *rep* gene block (Twist Biosciences) was generated with three silent mutations in codons N328, T239, and I330, and this gene was cloned into pITR-p5-rep2-cap9 using Gibson Assembly to create pITR-p5-rep2-cap9_bc. To create vectors for assessing DNA packaging of individual variants, the ITRs were removed from pITR-p5-cap9-rep2 with polymerase chain reaction (PCR) and cloned into the same backbone using Gibson Assembly to create pp5-cap9-rep2. Vectors for expressing Rep with synonymous mutations within the p19 promoter were constructed by cloning oligos from Twist Bioscience into pp5-cap9-rep2. Stop codons were generated in rep within pp5-cap9-rep2 using site-directed mutagenesis. Plasmids were verified by Sanger sequencing (Azenta) or whole-plasmid sequencing (Plasmidsaurus).

### AAV production for variant characterization

AAV vectors were produced using a double-plasmid polyethylenimine (PEI) transfection in HEK293T cells in 15-cm culture dishes. For these transfections, a mixture of DNA was used that included the pXX6 helper plasmid encoding adenoviral helper genes (19 µg), pRep2-Cap9, a rep2-cap9 mutant genome that lacks ITRs and thus cannot be packaged (10 ug), and pAAV-CMV-GFP, a plasmid encoding ITR-*gfp* that uses a CMV promoter to drive transcription (10 µg). After 24 hours, the cells were replenished with fresh serum-free media. After 48 hours, cells were harvested using a cell scraper. Cells were resuspended in 1 mL gradient buffer (10 mM MgCl2, 150 mM NaCl, 10 mM Tris, pH 7.6) and lysed using three freeze (with liquid nitrogen) and thaw cycles. The solution was pelleted by centrifugation (3000 x g for 10 min), and the supernatant was saved. Prior to virus quantification, this mixture was treated with Benzonase Nuclease (Sigma) for 40 minutes at 37°C to remove residual DNA to prevent unpackaged nucleic acids from being amplified during quantitative PCR (qPCR).

### Virus Quantification

Viral transgene titers were quantified using qPCR using primers provided in Table S2. Following Benzonase treatment, viral capsids were denatured in 2M NaOH at 56°C for 30 minutes to release encapsidated DNA. The mixture was neutralized using 2M HCl. The neutralized mixtures with viral DNA were diluted 1:100 in ultrapure water (Invitrogen) containing 10 ng/mL sheared salmon sperm DNA (Fisher Scientific). qPCR reactions were prepared by mixing this viral DNA solution (4 μL) with 2X SYBR Green PCR Master Mix (Life Technologies) (10 μL) and primers whose final concentrations were 0.2 μM. The total reaction volume was 20 μL. The qPCR thermal cycler protocol was: 95C for 10 minutes, 40 cycles of 95°C for 30 seconds, 55°C for 30 seconds, and 72°C for 30 seconds, with a stepwise increase from 40°C-95°C for 5 seconds each. Serially diluted recombinant AAV transgene (pITR-p5-cap9-rep2) or synthetic GFP flanked by ITRs (pITR-CMV-gfp-ITR) was used to generate a standard curve. Samples were analyzed using a BioRad CFX96 qPCR machine. Data analysis was performed on Microsoft Excel. Additional qPCR information is available in the Supplementary Information.

### Evaluating cross packaging

To minimize cross-packaging and genotype-phenotype mismatches, the amount of DNA used for transfections was analyzed. To do this, mixtures of pITR-rep2-cap9-ITR and pITR-CMV-gfp-ITR were transfected in parallel into HEK293T cells on 15 cm dishes (23). These transfections used varying amounts of pITR-rep2-cap9-ITR (10, 50, 500, and 5000 ng), pITR-CMV-gfp (10, 50, 500, and 5000 ng), and 19.99 ug pXX6. The latter vector was required as a helper vector as previously described (21). The cross-packaging assay was performed in the presence and absence of 10.01 ug herring DNA (Sigma). The herring DNA was included to even out the stoichiometry of DNA-PEI complexes. Cross-packaging was measured using qPCR. With each sample, reactions were performed using either a primer pair targeting the CMV promoter or a primer pair targeting wildtype *rep2*. The PCR cycling conditions were identical for each reaction. The qPCR thermal cycler protocol is the same as mentioned previously. This approach simulates a library transfection, but it is easier to quantify with only two types of cargo in the library. With this analysis, the signal from the CMV promoter analysis represents the cross-packaged DNA, while the signal from the *rep2* reaction represents viruses that correctly packaged an AAV genomes.

### Library synthesis

Synthetic DNA encoding all possible single amino acid changes in the Rep gene was computationally designed using Python scripts adapted from Coyote-Maestas (24). Oligonucleotide pools encoding each mutation were synthesized in tiles (227 to 231 base pairs) flanked by BsmBI restriction sites (Twist Biosciences). Primers were computationally designed to generate BsmBI restriction sites following PCR amplification of the oligo tiles and corresponding backbones using pITR-p5-cap9-rep2 as template. Figure S1 shows how each tile ensemble was cloned into the plasmid backbone using Golden Gate assembly (25). For each cloning reaction, tiles were PCR amplified while pITR-p5-cap9-rep2 was amplified as the vector backbone. Amplicons were gel purified using agarose gel electrophoresis and a Zymoclean Gel DNA Recovery Kit (Zymo Research). The PCR reactions using pITR-p5-cap9-rep2 as template were treated with Dpn1 (20 units) following the reaction to remove residual methylated parental DNA that could affect the cloning downstream. Amplicons were then purified using agarose electrophoresis and assembled using Golden Gate assembly (25). Enzymes for cloning were from New England Biolabs.

Each vector ensemble was electroporated into NEB Stable *E. coli,* plated on LB-agar medium containing carbenicillin (100 µg/mL), and incubated at 30°C for 24 hours. For each tile reaction, cells were spread on multiple LB-agar plates containing carbenicillin (100 μg/mL) to ensure adequate coverage of the library. In cases where transformation yielded less than 2000 colonies, the cloning process was repeated to increase coverage until a minimum of 2000 of CFUs were obtained for each tile. According to NGS, we obtained coverage of >90% of the total library, with Tile 1 presenting the lowest coverage, 72%; the proximity of this tile to the ITR is thought to have created the cloning challenges. To harvest each tile library, ∼2 mL of LB medium was added to each agar plate, a sterile spreader was used to pool the colonies, and the resulting slurry was transferred to 50 mL conical tubes. After mixing, the Zymo Research Plasmid Miniprep kit (transfection-grade) was used to isolate the plasmid ensemble encoding each tile library. This pool was sequenced for each tile prior to any further manipulations (Figure S2). Prior to transfections, the mutant library was mixed with the positive (pITR-p5-rep2-cap9_bc) and negative (pITR-p5-rep2-cap9_K340H) control vectors at ratios of 90:5:5. These controls were included to provide a frame of reference for native AAV and an inactive AAV.

### Library selection

To select for DNA packaging, PEI transfection was performed using HEK293T cells in 15 cm culture dishes. All selections were performed in triplicate. Tiles were subpooled in 5-plate transfections, such that one or two tiles were evaluated in each transfection. For each transfection, we used a DNA mixture containing pXX6 helper plasmid (19 µg) encoding the adenoviral helper genes, an ITR-rep2-cap9 plasmid mixture (20 ng) encoding the Rep2 point-mutant library and embedded control vectors, and sheared herring DNA (∼21 µg). After 48 hours, cells were harvested and lysed using three freeze-thaw cycles, cells were treated with 50 units/mL Benzonase Nuclease for 40 minutes at 37°C, and the mixture was layered onto a 15% to 54% iodixanol gradient as previously described (10). Tubes were ultracentrifuged at 48,000 rpm (169,021g) usiung a Beckman Type 70Ti rotor for 105 minutes at 18°C. Viruses were extracted from the 40% iodixanol layer with an 18-gauge needle and 3 mL syringe. Virus preps were stored at 4°C in 3 mL cryovials.

### Next-Generation Sequencing (NGS)

To evaluate the sequence diversity of the naive library, amplicons for short-read NGS across the rep gene were generated by PCR amplifying the individual tiles and adding Illumina adapters (Genewiz Amplicon-EZ). To assess sequence diversity following virus production, MiSeq (Illumina, Inc) was used to analyze the plasmid DNA used for transfections and DNA packaged by AAV following virion purification. A custom script was used to analyze the sequence diversity in each tile, which is provided in Table S3. The fitness for each mutant was evaluated with the following expression:

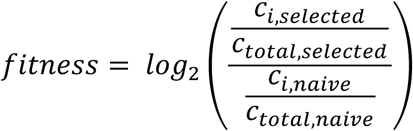

where *c* is the number of counts for either a mutant *i* or the total reads in the NGS run.

### Statistical Analysis

To evaluate the relative DNA packaging of Rep mutants containing a stop codon or synonymous mutations in the p19 promoter, we used single-factor ANOVA to assess statistical significance. We then used a post-hoc Tukey t-test to identify the groups with different means.

## Results

Cells produce AAV as a heterogeneous mixture, at times with more than half of the capsids lacking packaged DNA (4–6). This observation suggests that cellular selections could be used to mine libraries of AAV mutants for variants with improved DNA packaging (26). To first establish the extent to which virus production depends upon the Rep protein, we transfected HEK293 cells with the full-length AAV genome and a genome containing a K340H mutation in Rep (Figure 1A), which is deficient in helicase activity (27). For these studies, we used a chimera of the AAV2 and AAV9 genomes, which had Rep from AAV2 and Cap from AAV9. This genome was used because AAV2 Rep is frequently used for therapeutic rAAV production to package DNA cargo in AAV9 Cap, which is a capsid serotype used in approved gene therapies. Following virus production and purification, qPCR analysis revealed >3,600-fold decrease in viral genomes yield for the K340H Rep mutant compared with a native Rep (Figure 1B). This finding shows that the K340H mutation disrupts Rep activity and reveals that these vectors can be used as negative and positive controls when performing library selections.

**Figure 1.**
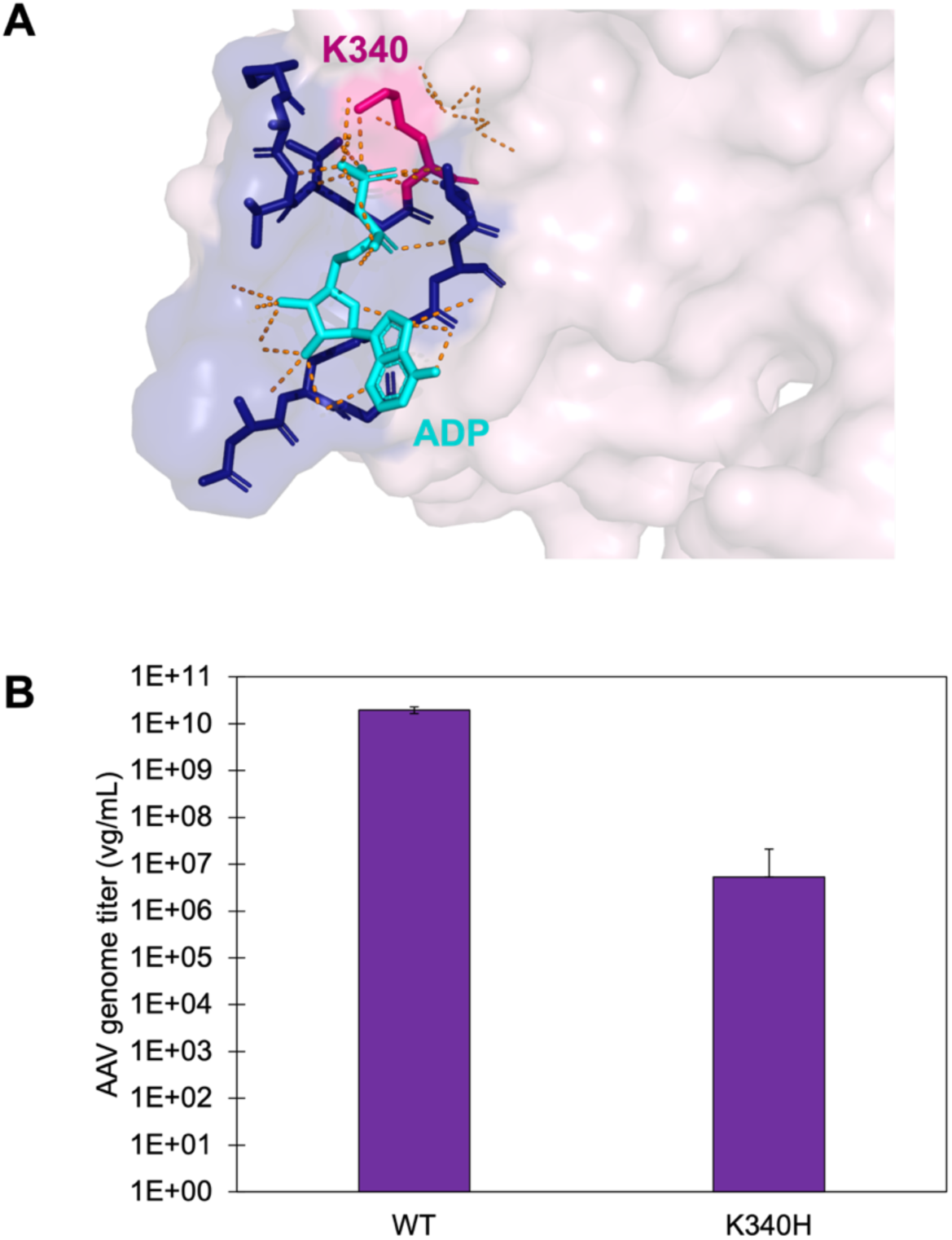
Transfection of AAV containing a K340H mutation in Rep. (**A**) To evaluate the activity of Rep, a K340H mutation was generated in the genome. Residue K340 directly interfaces with ATP in the helicase region and is required for ATP-dependent helicase activity. Dashed lines are noncovalent interactions. Image generated with Pymol. (**B**) Data obtained for transfections using the native genome (WT) and a genome harboring the K340H mutation. The data represents the average calculated from 2 biological replicates and 3 technical replicates, with error bars representing ±1 standard deviations. The viral genomes obtained for WT are significantly higher than K340H (ANOVA single-factor, p value = 0.003).

One challenge with selecting AAV mutant libraries for variants that alter virus production is the loss of genotype-phenotype linkages, which can occur under transfection conditions that favor cross-packaging (23, 28). To establish conditions that maintain genotype-phenotype linkages, we investigated the relationship between the amount of AAV genomic DNA transfected and the rate of cross-packaging. For these experiments (Figure 2A), we transfected equimolar mixtures of plasmids coding for the AAV genome and DNA encoding a GFP reporter (20 to 10,000 ng each), helper plasmid, and herring DNA to even out the stoichiometry of PEI to DNA. Following virus production and purification, qPCR was used to analyze the relative concentrations of AAV genome versus the GFP reporter that had been packaged. At the highest DNA transfection level (10,000 ng), most viruses were packaged with the GFP reporter in the absence (Figure 2B) of herring DNA. In contrast, at the lowest levels of DNA transfected (20 ng), only 8 ±4% of the viruses contained the GFP reporter. While this condition yielded the lowest cross packaging, it also presented lower total viral yields compared to the other conditions screened. When performing this experiment with herring DNA, we found that the transfections with herring DNA resulted in higher viral genome titers than the transfections without herring DNA (Figure S3). Again, the lowest level of DNA minimized cross packaging (Figure 2C). These results identify conditions where mixtures of AAV mutant genomes can be selected to maximize genotype-phenotype linkages and they reveal the importance of including herring DNA to maximize viral genome yields.

**Figure 2.**
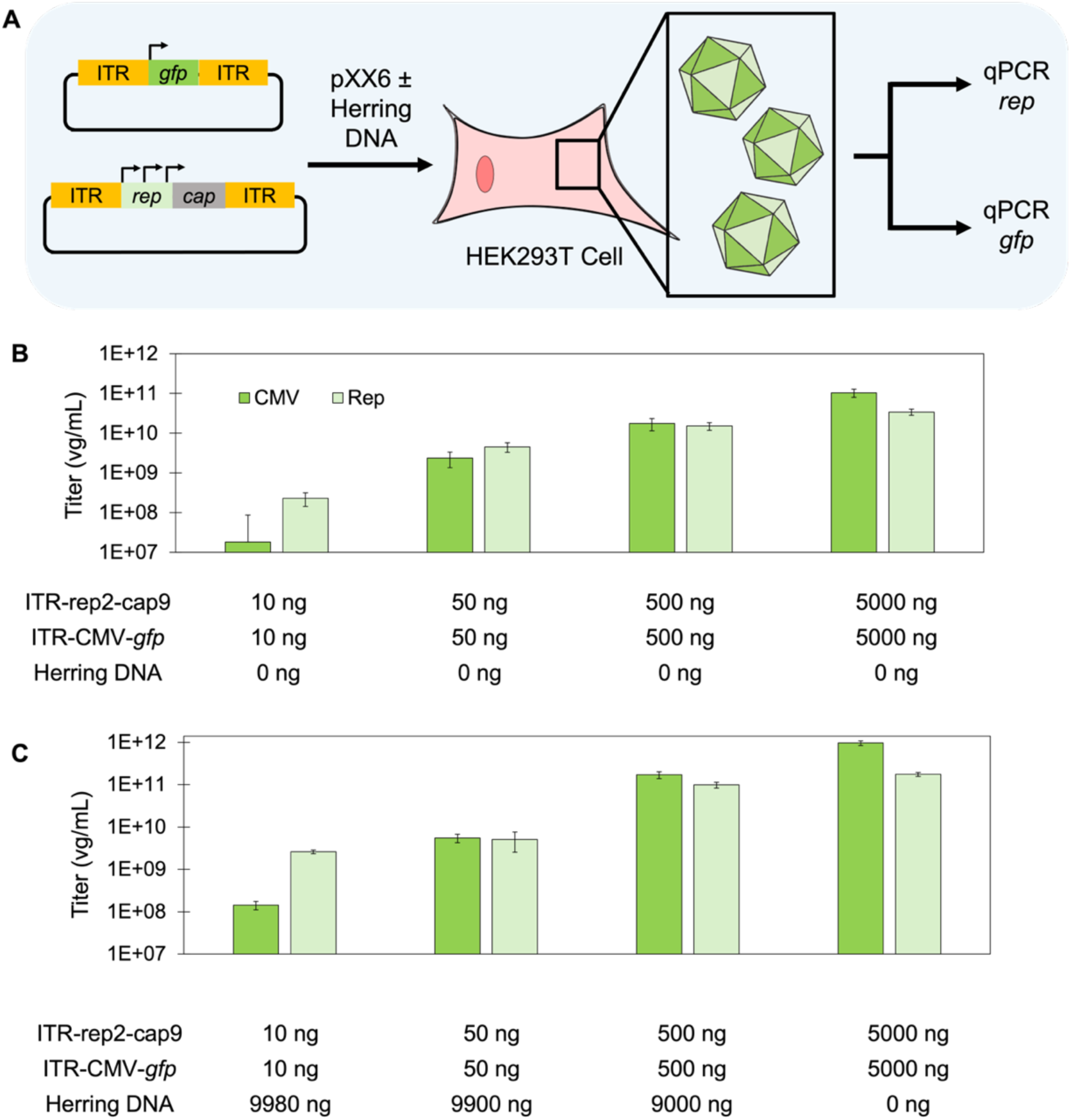
Minimizing cross packaging of DNA in capsids. (**A**) Cross packaging was evaluated by mixing ITR-CMV-gfp and ITR-rep2-cap9 prior to transfection into HEK293 cells. After harvesting virus that was produced, multiplexed qPCR was used to quantify each cargo by leveraging primers targeting the CMV promoter in ITR-CMV-gfp and *rep* in ITR-rep2-cap9. Experiments performed in (**B**) the absence or (**C**) presence of dummy DNA. For B and C, total virion formation having each DNA cargo was quantified. For each experiment, the data represents 4 biological and 3 technical replicates, with error bars representing ±1 standard deviation. When transfecting 20 ng of ITR-CMV-gfp and ITR-rep2-cap9 in the presence of herring DNA, the value obtained for ITR-rep2-cap9 was higher than ITR-CMV-gfp (two-sample t-test, p value = 0.12), presenting a ratio of ∼1:18 for CMV:*rep*. For the 20-ng transfection without herring DNA, the qPCR titer for ITR-rep2-cap9 was also higher than that of ITR-CMV-*gfp* (two-sample t-test, p value = 0.03), with a CMV:*rep* ratio of ∼1/12.

A library of Rep mutants was generated in the context of the AAV genome by synthesizing oligo pools encoding different amino acid substitutions and then inserting those into the genome using a tile based cloning approach (Figure S4A). Sequencing the vector pools for each tile revealed variation in sequence coverage, ranging from 72 to 95% of the expected mutations. Among all tiles created, 90% of all possible non-synonymous mutations were observed. On average, we sampled 17 different amino acid substitutions at each native position (Figure S4B), with gaps in coverage at the ends of each tile. Across the Rep gene, stop codons were observed at 94% of the native sites. These findings show that our tile-based cloning approach effectively samples the intended non-synonymous mutations as well as stop codons, which were intended to be coded in our library.

Our Rep mutant library was selected by performing transfections of HEK293T cells in replicate using pools of plasmids encoding the AAV genomes having non-synonymous mutations (Figure 3A). When transfecting low DNA concentrations that minimize cross packaging, virus yields are low. For this reason, selections were performed using plasmids containing one or two mutant tiles. In all cases, transfections included the parental AAV vector containing silent mutations for identification and a vector having the K340H mutation within the rep gene, which abolishes activity (27). After purifying viruses and isolating the ssDNA cargo, tiles harboring mutations were PCR amplified and analyzed using NGS. This analysis was performed on both the DNA used for transfection and selected libraries, and the resulting sequence counts from each were used to calculate enrichment scores for each mutant (Figure 3B). Among all of the amino acid substitutions sampled, prolines were the most consistently disruptive to Rep as observed in other protein engineering studies (9, 29). Through all experiments, the embedded positive control having three silent mutations and negative control containing the inactivating K340H mutation were observed, with the positive control presenting a baseline log fold enrichment of 0 and negative control being diluted by ∼100 fold (Figure S5). As the inactivating mutation is coded on Tile 6, we only used it to assess how well the experiments were performed rather than tying them to the NGS analysis.

**Figure 3.**
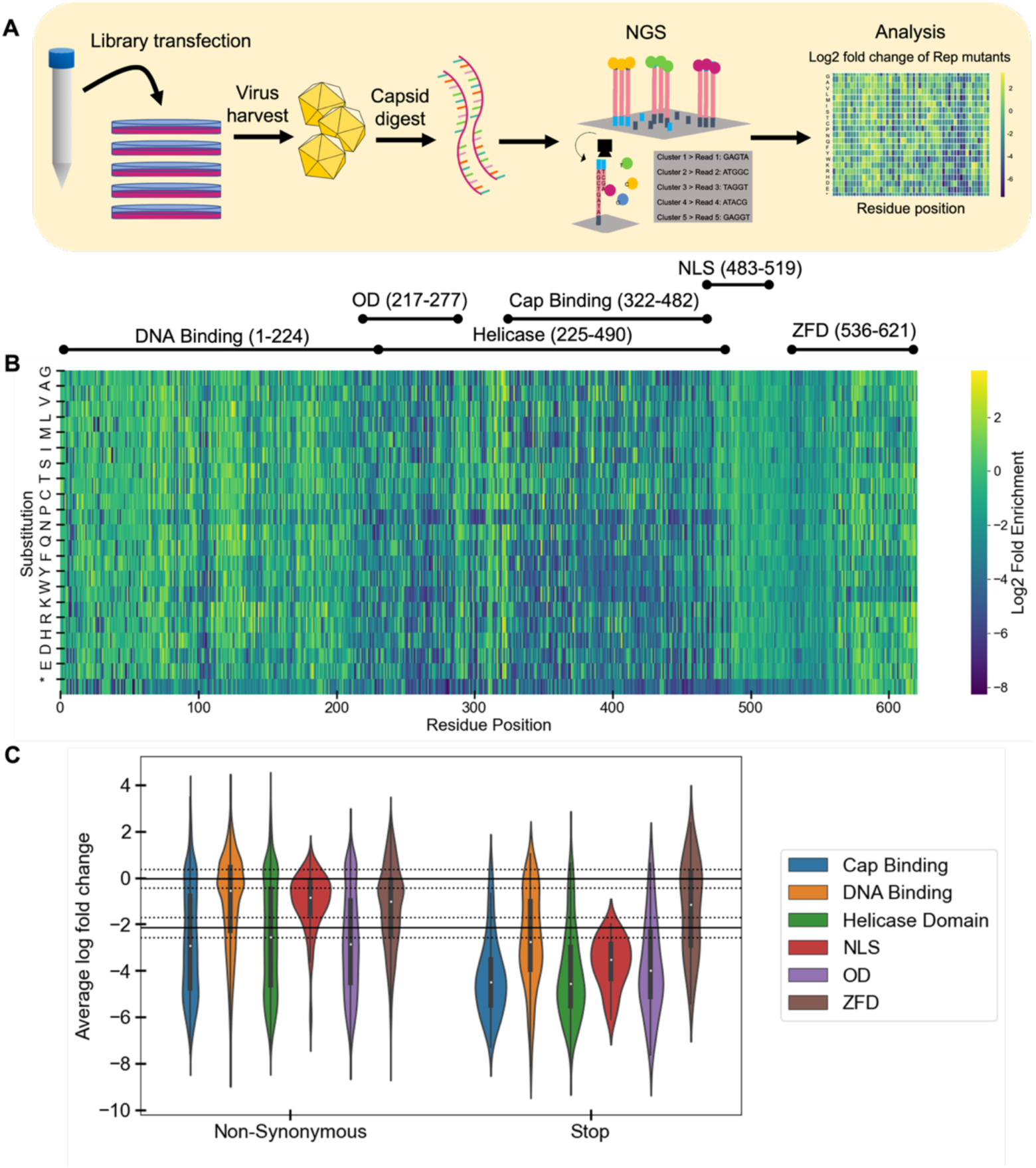
Non-synonymous mutant enrichment following selection. **(A)** To identify Rep with improved activities, mutants were transfected into HEK293 after mixing with the positive and negative controls at ratios of 18:1:1. This mixture (20 ng) was mixed with helper plasmid (XX6) and herring DNA prior to transfection. Each selection was performed triplicate. (**B**) The ratio of selected to non-selected DNA was calculated for each tile to establish the enrichment for each possible non-synonymous mutation. Values shown represent the average from the three biological replicate selections performed. The x-axis represents the residue mutated in the primary structure while the y axis represents the amino acid substitution at each position. Function regions of the protein are marked above the heatmap. (**C**) Violin plots enrichment show the dispersion of enrichment values across each domain for non-synonymous mutations and stop codons. Solid black lines represent the enrichment of the positive and negative controls. Dashed lines represent one standard deviation of the mean. NLS, nuclear localization signal; OD, oligomerization domain; ZFD, zinc-finger domain.

To assess how Rep mutational tolerance varies at the domain level, we compared the dispersion of fitness values in all domains (Figure 3C). When all mutations were analyzed for each domain, two major modes were observed, which were interpreted as representing functional and loss-of-function Rep variants. Mutants in the nuclear localization sequence, origin-binding domain, and zinc-finger domain presented the highest fraction of variants within the mode with the higher enrichment values. In contrast, mutations in the cap binding, helicase, and oligomerization domains all presented a greater dispersion of fitness values.

As the abundances of parental AAV following selection were being analyzed, we mined our sequence data for the most highly enriched sequences, regardless of whether they were purposefully designed in our sequence diversity. With the selection of tile 2, this analysis revealed that four of the ten most abundance sequences had synonymous mutations (Figure 4A). In contrast, when the naive library was analyzed, these mutants were present at trace levels (Figure S6). When the most highly enriched synonymous mutants were analyzed, we observed enrichment values that exceeded the positive control and all other non-synonymous mutations (Figure 4B). This finding suggests that synonymous mutations can modulate the DNA packaging activity of Rep, at least when the AAV genome represent the DNA cargo being packaged.

**Figure 4.**
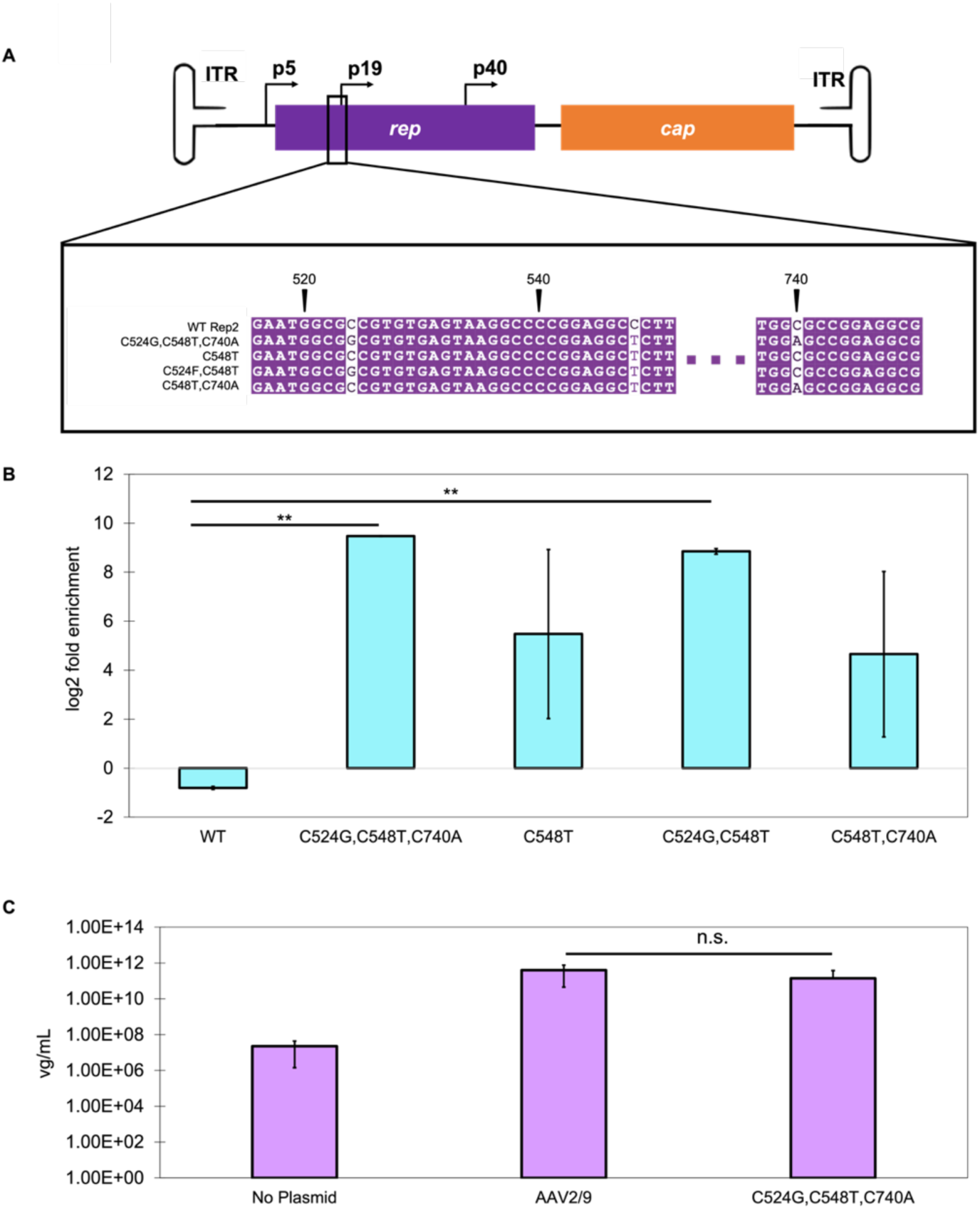
Synonymous mutations enriched by selections. (**A**) Silent mutations found near the p19 promoter have 1-3 nucleotide substitutions compared to WT Rep2. (**B**) The enrichment of AAV variants whose abundances increased dramatically following the selection. These variants had different combinations of synonymous mutations proximal to the p19 promoter. All of these variants presented enrichments that are significantly higher than that of positive control (WT) in the selection. (**C**) Individual virus preparations were generated for the positive control (WT) and the most enriched variant having synonymous mutations. Following packaging of ITR-CMV-gfp, the titer of synthetic cargo packaged was measured. The value obtained for the mutant and WT were not significantly different (**, p < 0.0001; n.s., not significant). The data represents 3 biological and 3 technical replicates, with error bars representing ±1 standard deviations.

To investigate how the enriched synonymous mutations impact Rep activity with a non-native cargo, we generated the synonymous mutations (C524G, C548T, C740A) that were enriched to the greatest extent in a vector containing the rep-cap genes but lacking the ITRs. To study the DNA packaging activity, we transfected this vector in parallel with the helper plasmid (pXX6) and a vector containing ITR-*gfp* under transcriptional control of a CMV promoter. Viruses were then purified, and qPCR was used to quantify the amount of packaged DNA (ITR-*gfp*) using primers that target the CMV promoter (Figure 4C). The variant having synonymous mutations near the p19 region of Rep presented viral titers that were similar to the native Rep construct; these values were not significantly different from native rep (p = 0.21; ANOVA single-factor). Thus, a Rep having the synonymous mutations enriched in library selections packages capsids with synthetic DNA cargo (ITR-gfp), but the efficiency of this packaging is not improved compared to a native Rep protein.

## Discussion

The most striking mutations enriched by our selection were synonymous mutations proximal to the p19 promoter. The four AAV variants identified with high enrichment resulted from proximal sequence changes within the Rep coding sequence, including: (i) C524G, C548T, and C740A, (ii) C548T, (iii) C524G and C548T, and (iv) C548T and C740A. These mutants all presented low abundances in the naive library before selection, representing 0 to 0.01% of total counts in their respective tiles. Following selection, they all presented 10^2^ to 10^4^-fold increases in abundance. Two of these mutants achieved log fold enrichments that were statistically significant compared to wildtype (p = 8.96E-10 and p = 2.14E-08 for mutants C524G/C548T/C740A and C524G/C548T, respectively). This trend can be contrasted with our positive control, whose abundances remained similar before and after the selection, and the negative control which was consistently diluted by the selection. The p19 promoter is responsible for transcribing Rep52 and Rep40 (30), which play a role in DNA cargo packing (17, 18). The p19 promoter is also activated by the Rep78 and Rep68 proteins (31), implicating these mutations in modulating protein-nucleic acid interactions critical to DNA packaging.

Activation of the p19 promoter relies on Rep-Binding Elements (RBEs) within the p5 promoter and/or ITRs (31). Of note, the RBEs possess several GCTC motifs (32), which are thought to mediate protein-DNA interactions (32–34). GCTC motifs are also present on the ITRs and are required for Rep binding and packaging (32–34). When no ITRs are present, p19 activation depends on the p5 RBE and p19 Sp1 site (31). The proposed role of the p5 RBE and p19 Sp1 sites is to act as a scaffold for bringing transcription complexes from p5 closer to the p19 promoter, which leads to the initiation of DNA replication and expression (31, 35). Rep78/68 also covalently binds to the RBE on the ITR and nicks the ITR at a sequence called the terminal resolution site (trs) to initiate genome replication (36). Interestingly, the p19 mutants enriched to the greatest extent in our selection generated 1 to 2 additional GCTC motifs. Rep can also bind variations of this motif with weaker affinity, and it is hypothesized that this motif may regulate dynamic Rep-DNA interactions (32). How this mutation affects protein-DNA interactions is not known. It is possible that this sequence change affects Rep-DNA, TF-DNA, Sp1-DNA, and/or cAAP interactions (35).

When packaging the synthetic GFP transgene DNA cargo, mutation of the p19 promoter did not lead to enhanced packing, although it presented activity that could not be differentiated from the native AAV. This observation can be contrasted with the results from our selection, where large enrichments were observed. These differences could arise because the synthetic cargo (ITR-CMV-gfp) packaged for the individual mutant virus preps does not contain a p19 promoter or GCTC motifs like the native genome (ITR-Rep-Cap). In the future, it will be interesting to explore how other silent mutations proximal to this promoter affect packaging efficiency and whether GCTC-like motifs can be identified that could be introduced to synthetic cargos to enhance interactions with Rep and increase synthetic cargo packaging efficiency in a similar manner.

Our analysis of Rep tolerance to non-synonymous mutations mirrors a recent study that explored mutational tolerance across multiple AAV serotypes (19). In both studies, transfections served as the selection for Rep activity, and the incidence of cross-packaging was minimized. However, these studies differed in the strategy used to detect mutations. Herein, we sequenced the Rep gene directly, while the prior study used barcodes amended to the genome, which do not report on mutations generated during library synthesis, propagation, and selection. Our results suggest that future studies should use long-read sequencing to analyze AAV libraries following selection as this approach would provide insight into the abundances of intended mutations, mutations generated during library generation and manipulation, and the linkage of different mutations across the full AAV genome.

In our selections, a subset of the stop codons across the different Rep domains were enriched to a greater extent than the embedded negative control, implicating these variants as catalytically active. In a prior study, Rep was found to retain activity following truncation within the ZFD, showing that this domain is not necessary for production of AAV upon transfection, although it may still be necessary for some Rep functions (6, 19). However, truncations removing portions of the OBD and HD are not expected to be competent for DNA packaging as these domains are essential for DNA binding and helicase activity (15). Whether the Rep variants having stop codons in these domains translate functional protein will require future *in vitro* studies of helicase activity. RNA viruses can experience stop codon readthrough at detectable frequencies to expand the coding space of their minimized genomes (37). Stop codon readthrough rates also depend on the nucleotide context and oxidative stress (38–41), which could lead to the variability in stop codon read through observed. Additionally, the use of PEI for transfection may contribute to this trend as PEI can stimulate oxidative stress in cells (42), which has been shown to increase stop codon readthrough rates (40).

## Data Availability

The three most abundant nonsynonymous p19 *rep* mutants were deposited in GenBank. The accession numbers are PQ515803, PQ515804, and PQ515805. Raw NGS fastq files are available in NCBI’s Sequence Read Archive via submission number SUB14813973. Code for generating mutant libraries and analyzing NGS data are available on FigShare: DOI, <u>10.6084/m9.figshare.27328350.v1</u>, <u>m9.figshare.27328353.v1</u>, and <u>m9.figshare.27328404.v1</u>. qPCR data following the MIQE guidelines are available in the Supplementary Data, including standard curves (Figures S7 to S10) and no template control results (Tables S4 and S5), and the Materials and Methods.

## Supplementary Data

Supplementary Data are available at NAR Online.

## Funding

This research was supported by a sponsored research agreement supported by Biogen. In addition, TA was supported by NSF Research Traineeship grant 1828869 (to J.J.S.). D.M. is currently supported by the National Science Foundation Graduate Research Fellowship Program.

## Conflict of interest

The authors have no conflicts of interest to declare.

## Notes

### Competing Interest Statement

The authors have declared no competing interest.

### Summary of Updates

Very minor text edits were made to clarify the manuscript throughout.

