## Supporting Information for "Synonymous mutations in AAV Rep enhance genome packaging in a library selection"

**Table S1. List of plasmids.** For each plasmid, a brief description is provided as well as the origin of replication and the selection marker. CarR indicates carbencillin resistant, while AmpR indicates ampicillin resistant.

| Plasmid Name | Description | Origin of Replication | Selection marker |
|---|---|---|---|
| pITR-p5-rep2-cap9 | full AAV2 genome having Cap from AAV9 | pUC19 | AmpR, CarbR |
| pITR-p5-rep2-cap9_K340H | K340H mutation in Rep within ITR-p5-rep2-cap9 | pUC19 | AmpR, CarbR |
| pXX6 | helper plasmid containing adenoviral genes | pUC19 | AmpR, CarbR |
| pRep2-Cap9 | AAV2 genome having Cap from AAV9 but lacking ITRs | pUC19 | AmpR, CarbR |
| pAAV-CMV-GFP | CMV-GFP reporter flanked by AAV ITRs | pUC19 | AmpR, CarbR |
| p5-rep2-cap9_mut1 | C548T mutations within *rep* gene of pRep2Cap9 | pUC19 | AmpR, CarbR |
| p5-rep2-cap9_mut2 | C524G,C548T,C740A mutations within *rep* gene of pRep2Cap9 | pUC19 | AmpR, CarbR |
| p5-rep2-cap9_E205* | E205* mutation in Rep within pRep2Cap9 | pUC19 | AmpR, CarbR |
| p5-rep2-cap9_Y311* | Y311* mutation in Rep within Rep2Cap9 | pUC19 | AmpR, CarbR |
| p5-rep2-cap9_A527* | A527* mutation in Rep within pRep2Cap9 | pUC19 | AmpR, CarbR |
| pITR-p5-rep2-cap9_bc | full AAV2 genome with Cap from AAV9 and 3 silent mutations on Rep | pUC19 | AmpR, CarbR |

**Table S2. Primers used for qPCR.** The CMV primers are designed to amplify a 104 base pair region of the CMV promoter within pAAV-CMV-GFP, while the Rep primers are designed to amplify a 74 base pair region of the Rep gene.

| Name | Sequence |
|---|---|
| CMV-fwd | TCACGGGGATTTCCAAGTCTC |
| CMV-rev | AATGGGGCGGAGTTGTTACGA |
| Rep-fwd | CCGACTTTGCCAAACTGGTTC |
| Rep-rev | TCATCCACCACCTTGTTCCC |

**Table S3. Tile sequence diversity.** Following NGS analysis of each tile library, the fraction of single amino acids observed was quantified. In total, 11,160 of 12,400 possible mutations were observed with this sequencing, which represents 90% of the expected sequence diversity.

| Tile | Residues in Tile | % mutations observed |
|---|---|---|
| 1 | 1-62 | 71.7 |
| 2 | 63-125 | 93.7 |
| 3 | 126-186 | 93.4 |
| 4 | 187-249 | 85.8 |
| 5 | 250-310 | 95.1 |
| 6 | 311-372 | 90.9 |
| 7 | 373-435 | 92.7 |
| 8 | 436-496 | 93.9 |
| 9 | 497-558 | 92.9 |
| 10 | 559-621 | 93.5 |

**Table S4. NTC results for qPCRs for Figure 2, no herring DNA conditions.** The primers targeted CMV or *rep.* n=3 for each standard/replicate.

| **Standard / Replicate** | **Average Cq** | **Stddev Cq** |
|---|---|---|
| CMV / 1 | 35.88 | 1.17 |
| *rep* / 1 | 35.10 | 1.29 |
| CMV / 2 | 44.88 | 2.74 |
| *rep* / 2 | 35.28 | 0.54 |
| CMV / 3 | 40.42 | 2.46 |
| *rep* / 3 | 35.27 | 1.69 |

**Table S5. NTC results for qPCRs for Figure 2, herring DNA conditions.** The primers targeted CMV or *rep.* n=3 for each standard/replicate.

| Standard / Replicate | Average Cq | Stddev Cq |
|---|---|---|
| CMV / 1 | 39.25 | 1.58 |
| *rep* / 1 | 33.08 | 0.74 |
| CMV / 2 | 37.76 | 6.16 |
| *rep* / 2 | 33.17 | 1.58 |
| CMV / 3 | 37.92 | 1.65 |
| *rep* / 3 | 31.66 | 0.45 |

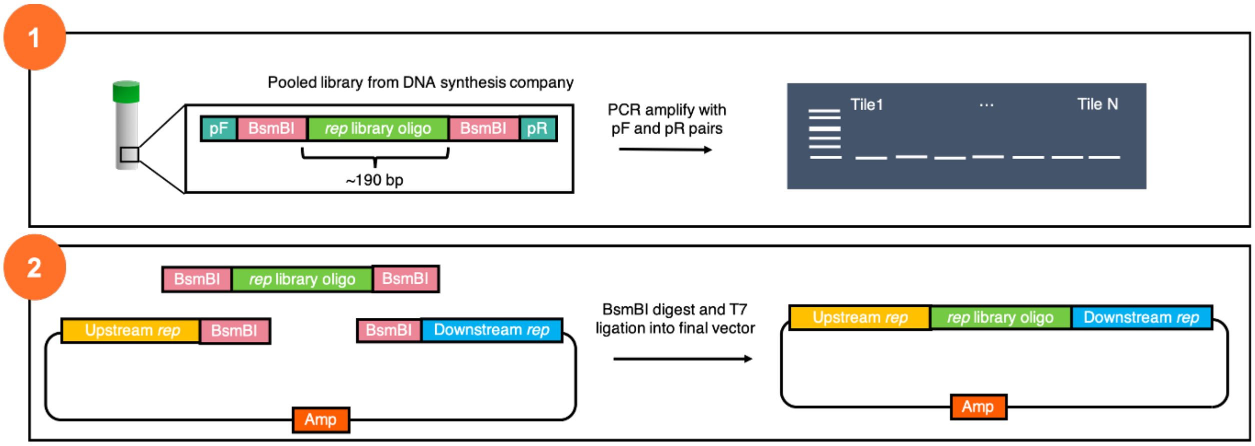


**Figure S1. Tile based cloning approach used for library construction.** First, we used software to divide the Rep gene into ten tiles having approximately 62-63 codons each; single-stranded DNA encoding every single amino acid substitution within each tile was designed computationally. The ten different oligonucleotide pools were commercially synthesized. Each oligonucleotide pool was amplified separately and cloned into pITR-p5-rep2-cap9. Following transformation into cells and purification of each tile's plasmid ensemble, next generation sequencing was performed to assess whether coverage was sufficient to proceed with selections. In cases where it was low, cloning was repeated.

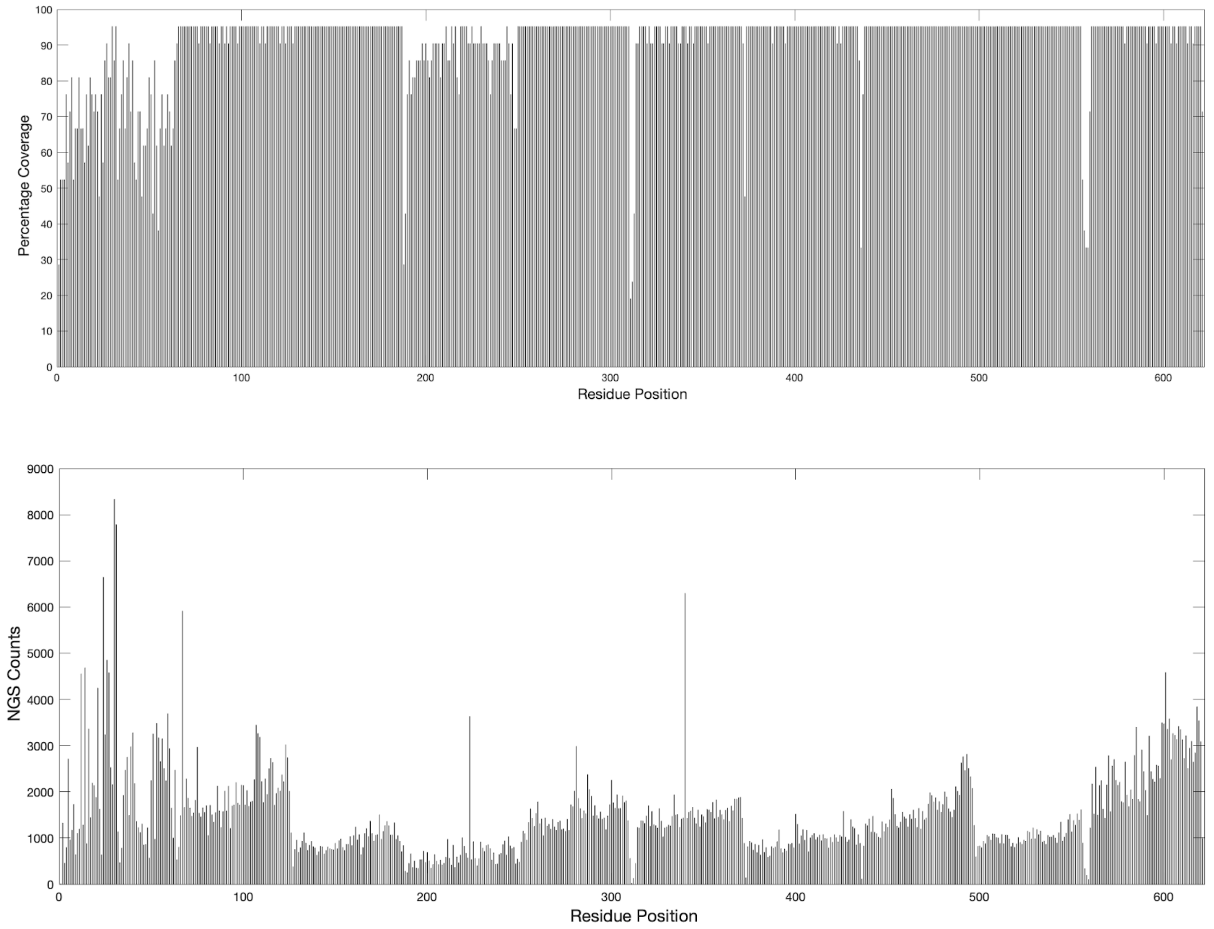


**Figure S2. Comparison of naïve library sequence coverage and depth of sequencing.** (**A**) The percentage of possible amino acids observed at each Rep codon with next generation sequencing of the individual tiles is compared with (**B**) the number of sequencing reads that observed a mutation at each native position.

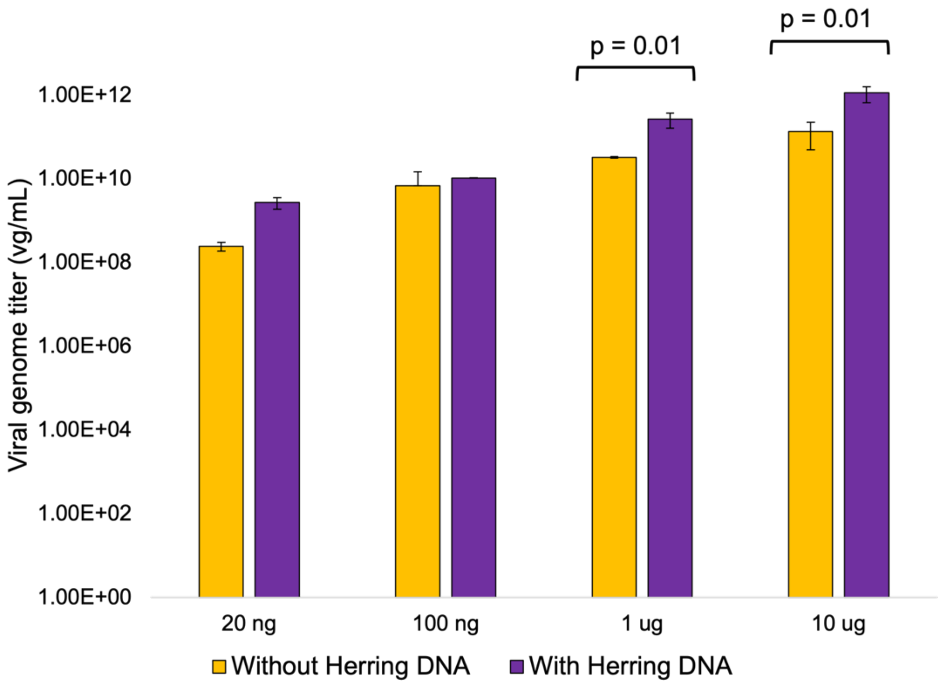


**Figure S3. Herring DNA increases the amount of total packaged DNA detected by qPCR.** The cross packaging assay was performed in the presence (purple) and absence (yellow) of herring DNA. For each condition, the total viral genome titer is shown, which represents the sum of the qPCR signals for virions that package the GFP reporter versus the AAV genome. The latter was quantified by analyzing *rep* copy numbers using qPCR, while the former was quantified using the copy number of the CMV promoter. p values were calculated using the two-tailed paired T-test. For 20 ng, 100 ng, 1000 ng, and 10,000 ng, respectively, p-values were 0.19, 0.47, 0.01, and 0.01.

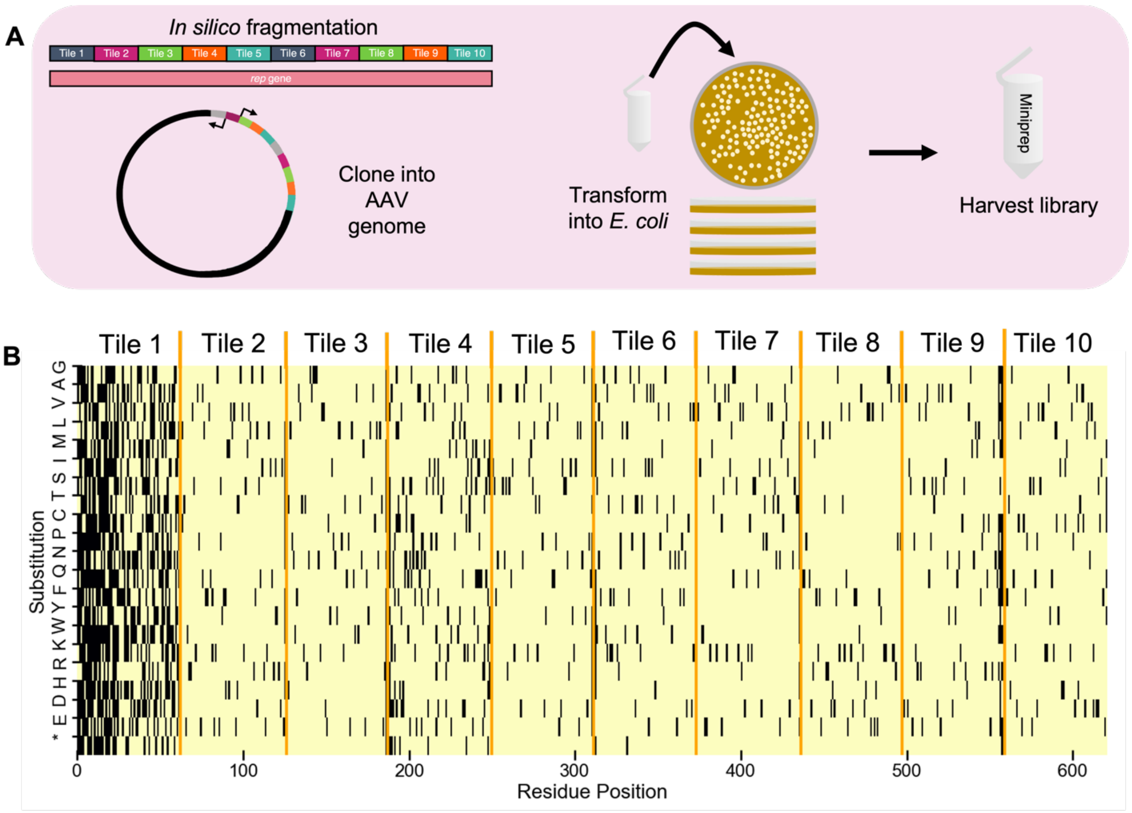


**Figure S4. Diversity of mutations sampled in the rep gene.** (**A**) Strategy used to construct and analyze the diversity found in the library. Computationally, the Rep gene sequence was fragmented into ten tiles *in silico*, each having a length (186 to 195 base pairs) that could be commercially synthesized using pooled oligonucleotide synthesis. The algorithm designed oligonucleotides encoding every unique non-synonymous mutation, and it added unique barcodes to the ends of each tile ensemble to allow for selective amplification. PCR amplified tiles were cloned into vector backbones complementary to each tile using Golden Gate cloning to generate an insertion library. (**B**) Deep sequencing was following the cloning of each tile to determine the non-synonymous mutations sampled. The initial library diversity before selection shows that we were able to generate 90% of the intended library. On the binary heatmap, pale yellow spots represent mutations that were present in the library and black spots represent mutations that were not in the naïve library. Each tile boundary is noted in orange.

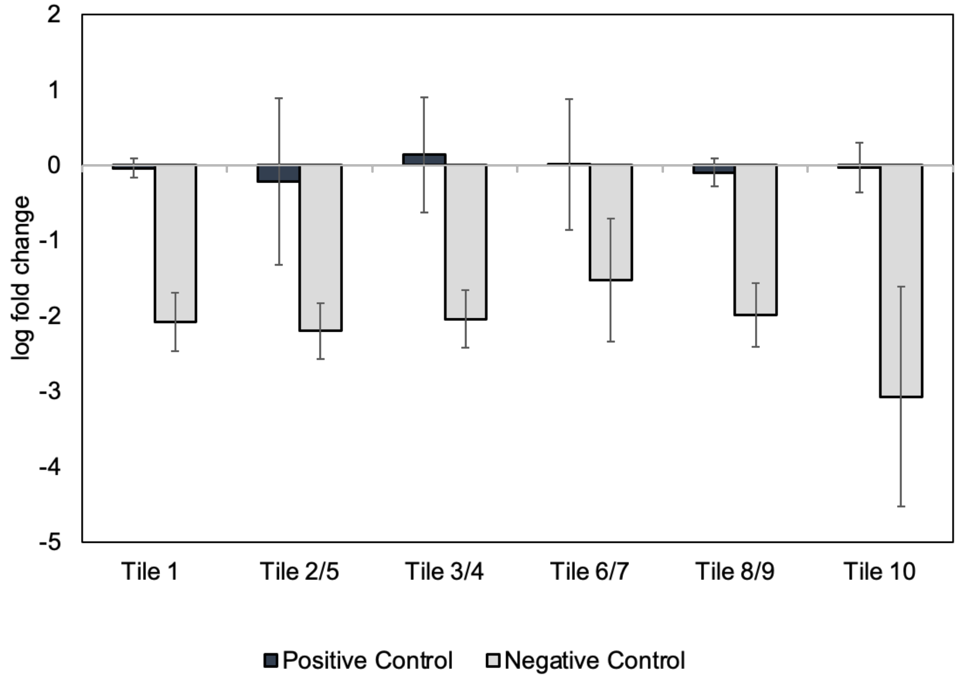


**Figure S5. Enrichments of controls vectors following library selections.** In total, six library selections were performed using mixtures of no more than two tiles and vectors encoding native Rep with silent mutations (pITR-p5-rep2-cap9) and Rep with an inactivating K340H mutation (pITR-p5-rep2-cap9_K340H). Each of these were performed in triplicate. The data shown represents the change in relative abundances of these control vectors for each individual biological replicate. Across all experiments, the abundance of the embedded positive control presented very small changes, while the negative control was diluted two to four orders of magnitude.

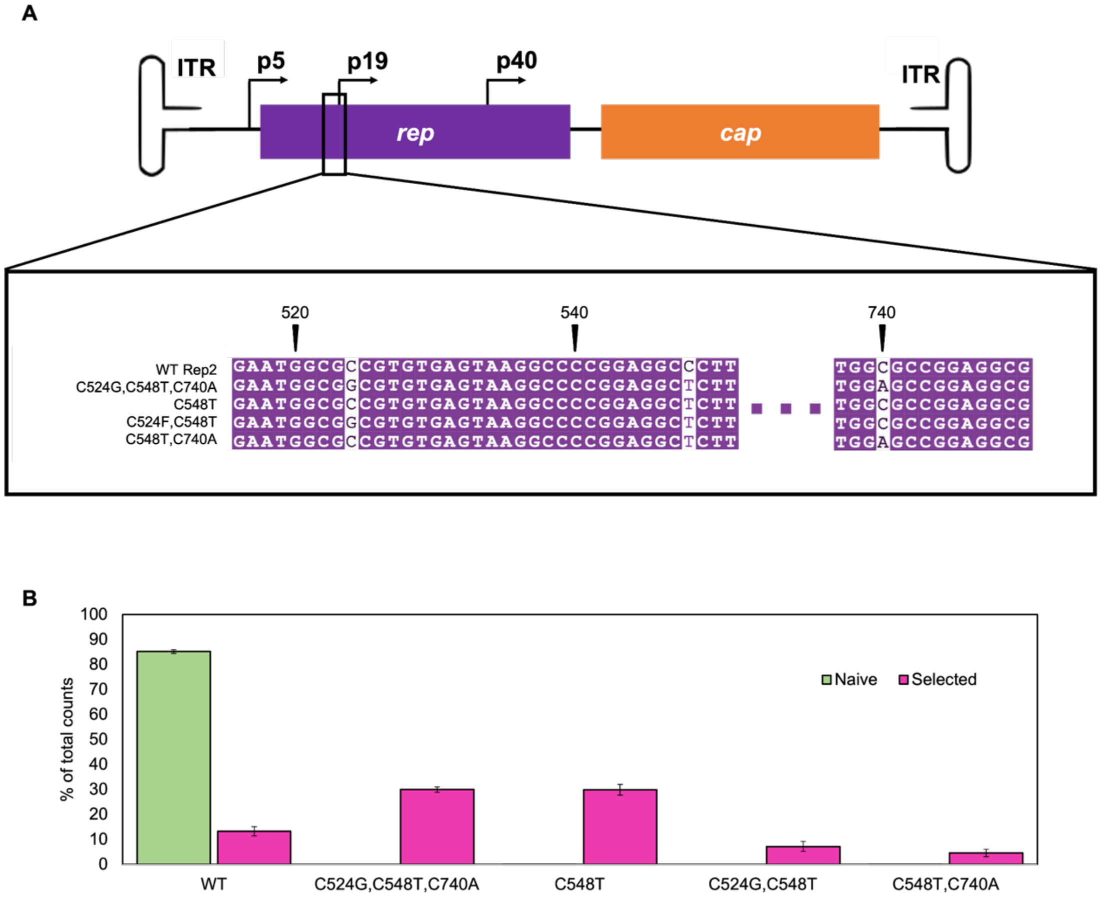


**Figure S6. Silent p19-proximal mutations within the AAV genome.** (**A**) Silent mutations found near the p19 promoter have 1-3 nucleotide substitutions compared to WT Rep2. (**B**) The percentage of NGS reads before and after selection for silent mutants increased while the percentage of reads before and after selection for WT decreased.

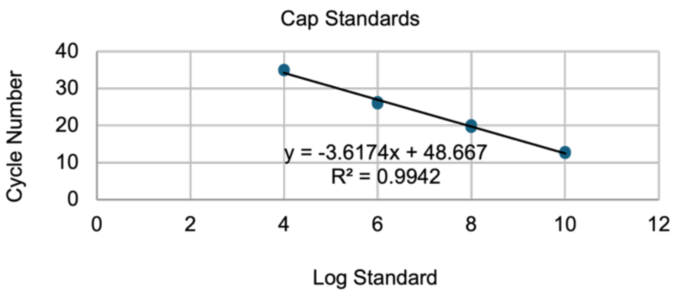


**Figure S7. qPCR standard curves for Figure 1 with primers targeting *cap*.** NTC results: average Cq = 32.35, stdev = 0.63, n = 3.

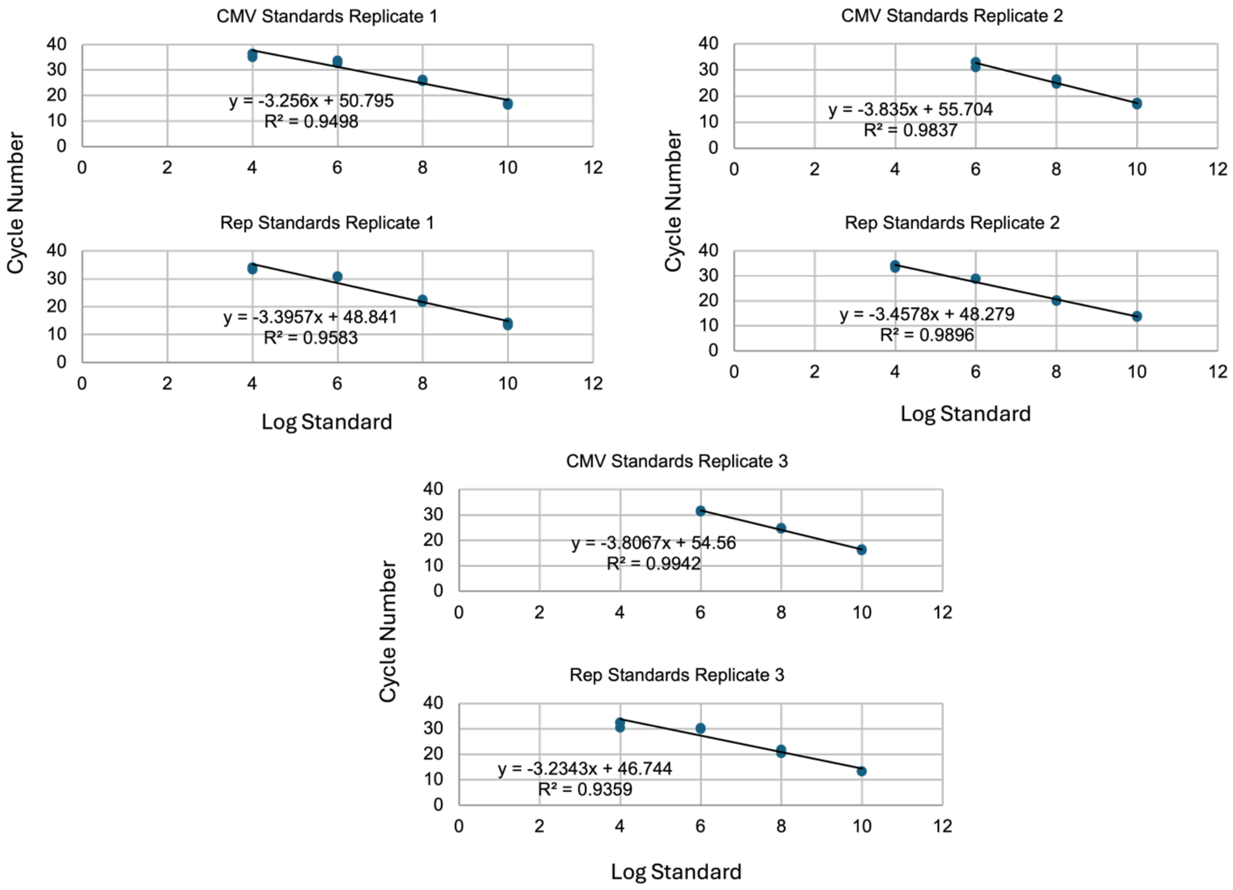


**Figure S8. qPCR standard curves for Figure 2, no herring DNA condition.** The primers targeted CMV or *rep*.

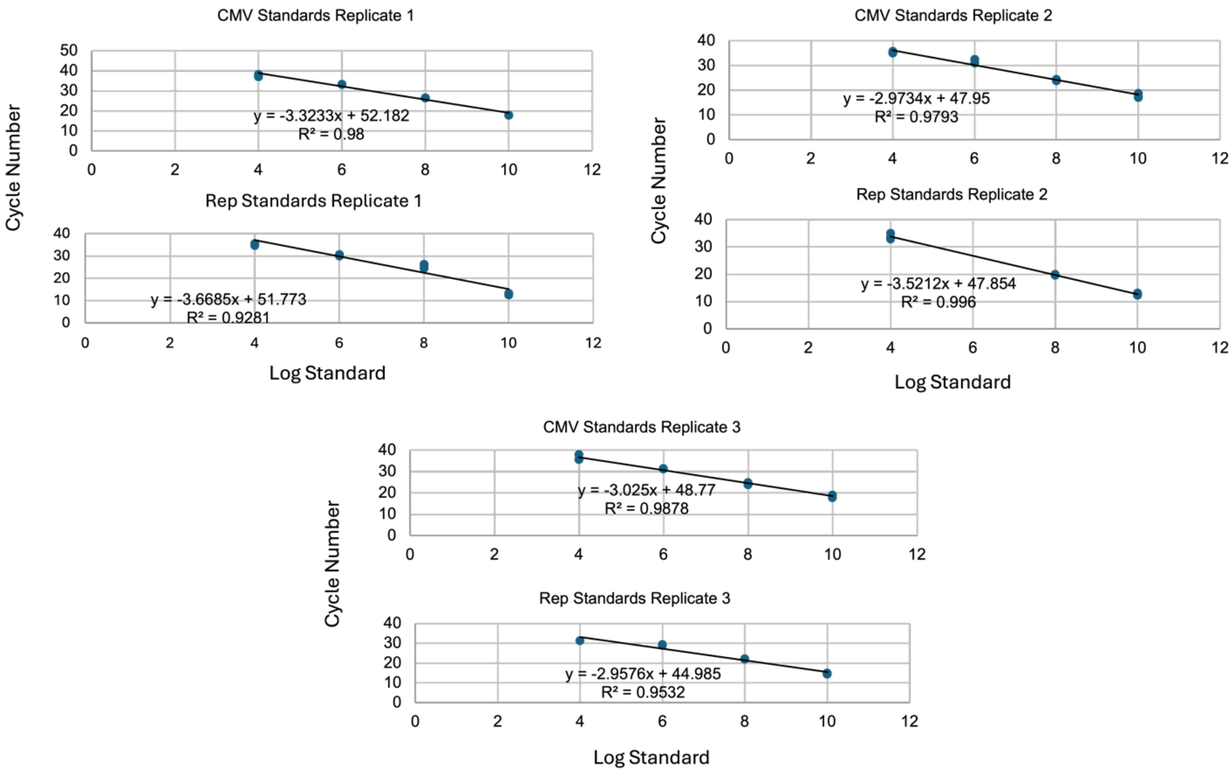


**Figure S9. qPCR standard curves for Figure 2, herring DNA condition.** The primers targeted CMV or *rep*.

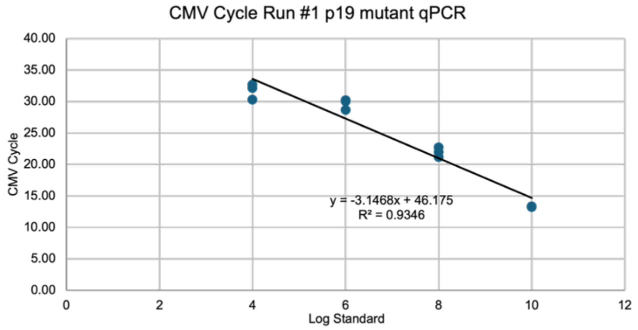


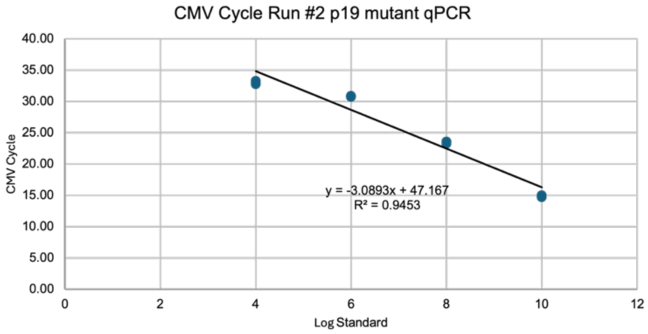


**Figure S10. qPCR standard curves for Figure 4 with primers targeting CMV.** NTC results for Run #1: average Cq = 34.93, stdev = 0.86, n = 3. NTC results for Run #2: average Cq = 35,89, stdev = 0.91, n = 3.
